# An inverse agonist of orphan receptor GPR61 reveals a novel allosteric mechanism

**DOI:** 10.1101/2023.05.01.538732

**Authors:** Joshua A. Lees, João M. Dias, Francis Rajamohan, Jean-Philippe Fortin, Rebecca O’Connor, Jimmy X. Kong, Emily Hughes, Ethan L. Fisher, Jamison B. Tuttle, Gabrielle Lovett, Bethany L. Kormos, Rayomand J. Unwalla, Lei Zhang, Anne-Marie Dechert Schmidt, Dahui Zhou, Kimberly A. Stevens, Kimberly F. Fennell, Alison E. Varghese, Andrew Maxwell, Emmaline E. Cote, Yuan Zhang, Seungil Han

**Affiliations:** Discovery Sciences, Medicine Design, Pfizer Inc., Groton, CT, USA; Internal Medicine Research Unit, Pfizer Inc., Cambridge, MA, USA; Internal Medicine, Medicine Design, Pfizer Inc., Groton, CT, USA; Internal Medicine, Medicine Design, Pfizer Inc., Cambridge, MA, USA

## Abstract

GPR61 is a biogenic amine receptor-related orphan GPCR associated with phenotypes relating to appetite and thus, is of interest as a druggable target to treat disorders of metabolism and body weight, such as obesity and cachexia. To date, lack of structural information or a known biological ligand or tool compound has hindered comprehensive efforts to study its structure and function. Here, we report the first ever structural characterization of GPR61, in both its active-like complex with heterotrimeric G protein and in its inactive state. Moreover, we report the discovery of a potent and selective small-molecule inverse agonist against GPR61 and structural elucidation of its unprecedented allosteric site and mode of action. These findings offer key mechanistic insights into an orphan GPCR, while providing both a new structural framework and tool compound to support further studies of GPR61 function and modulation.

## Main Text

### Introduction

G protein-coupled receptors (GPCRs) form one of the largest and most important classes of therapeutically relevant proteins in humans, accounting for an estimated 30-50% of the targets of currently marketed drugs^1, 2^. Nevertheless, a sizable fraction of disease-relevant GPCRs, particularly among orphan receptors, have not yet been harnessed for therapeutic modulation, though many are the focus of active research. GPR61, an orphan class A (rhodopsin family) receptor closely related to biogenic amine receptors^3, 4^, is predominantly expressed in the pituitary and appetite-regulating centers of the hypothalamus and brainstem. Mutagenesis and human genome-wide association studies have linked GPR61 to phenotypes associated with type 2 diabetes and body mass index^5,6^, making it a potential target for the modulation of appetite and body weight.

Like many other class A GPCRs^7^, GPR61 is a constitutively active receptor. It signals through Gαs to activate production of the small molecule second messenger cyclic AMP (cAMP) by adenylyl cyclase^8^. The mechanism underlying its constitutive activation remains poorly understood, but mutagenesis studies have suggested that residues near its N-terminus may play a key role^9^. A lack of structural information for GPR61 has been a significant impediment to progress in characterizing this receptor, in part because the absence of a known biological ligand or tool compound has made structural efforts challenging. Though 5-nonyloxy-tryptamine (5-NOT) has been reported as a GPR61 inverse agonist^10,11^, its low potency, low solubility, and lack of selectivity have limited its utility for pharmacological or structural studies. GPCR structural determination in the absence of a ligand or tool compound is fraught with challenges, including poor protein expression and solubility, often compounded by inherent structural plasticity that can be prohibitive for high-resolution structures. This is reflected in a relative paucity of unliganded GPCR structures in the Protein Data Bank. To provide insights into the structural basis of its constitutive activation, we report here the first unliganded structure of GPR61 in its active, G protein-coupled state, using cryo-electron microscopy (cryo-EM).

GPR61 knockout mice exhibit a hyperphagic phenotype leading to obesity^12^, suggesting that GPR61 inhibition by an inverse agonist might be used to treat wasting disorders, such as cachexia. As part of efforts to develop a small molecule therapeutic targeting GPR61 for the treatment of cachexia, we have discovered a sulfonamide inverse agonist tool compound that exhibits potent inhibition of GPR61 constitutive activity. Structural co-elucidation of GPR61 with this compound revealed it to act through a novel allosteric pocket by an unprecedented mechanism, binding and remodeling an intracellular pocket co-extensive with that normally occupied by Gαs in the activated state to block G protein activation. Collectively, the discoveries reported here shed light on the mechanism of GPR61 activation and a novel mechanism of GPCR inactivation, while providing a structural platform for future studies of GPR61 and a tool compound to support future mechanistic studies and drug discovery efforts.

## Results

### The structure of GPR61 in its active state suggests a basis for its constitutive activity

To better understand the molecular determinants of constitutive GPR61 activation, we determined its structure in complex with the Gαs/β1/γ heterotrimeric G protein by cryo-EM. While native complex assembly by co-expression of GPR61, Gαs, Gβ1, and Gγ was unsuccessful, fusion of a novel dominant negative^13^ variant of the Gαs/iN18 chimera^14^ to the receptor’s C terminus, combined with the use of a single-chain Fab (scFv16)^15^, stabilized formation of the GPR61-G protein complex. The Gαs/iN18 chimera forms, with Gβ, an epitope for scFv16, which stabilizes interactions between Gα and Gβ, while dominant negative Gα subunit variants have been used successfully to enhance the formation of GPCR-Gα complexes^13,16,17^. The structure of this stabilized complex was determined by cryo-EM at 3.5 Å nominal resolution (Fig. 1a). Each of the components of the complex was clearly resolved, with an overall architecture consistent with that of related active-state class A receptor structures^14,18–20^, suggesting the receptor was captured in an active-like conformation, though the binding of an agonist ligand may induce additional conformational changes.

**Figure 1.**
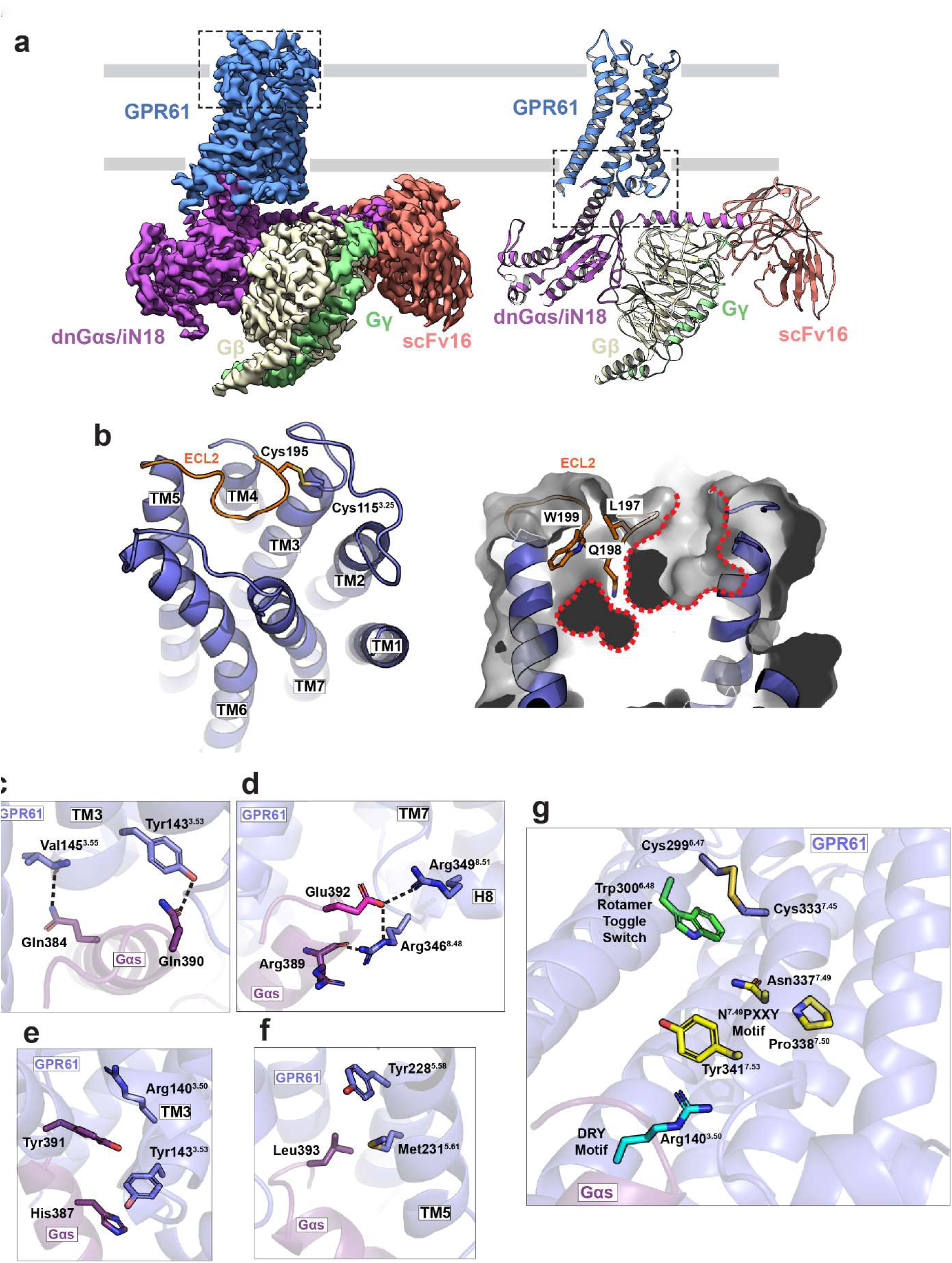
Cryo-EM structure of active-state GPR61-G protein complex. a. Map (*left*) and model (*right*) of active-state GPR61-G protein complex with scFv16, colored by subunit. Grey lines indicate the position of the plasma membrane. b. Highlighted by inset from A, left panel. *Left panel*, Extracellular face of GPR61, with ECL2 highlighted in orange and residues participating in the conserved disulfide shown in stick representation. *Right panel*, The orthosteric pocket of GPR61 is highlighted with a red dotted line. Key residues of ECL2 (orange) blocking access to the lower portion of the pocket are shown in stick representation. c-f. Key residues involved in GPR61-Gαs interaction, corresponding to the highlighted region of A, right panel. c. Polar interactions d. Hydrogen bond network underlying selectivity for Gαs. e. D/ERY motif hydrophobic stacking interactions with Gαs. f. Hydrophobic interactions. g. A novel disulfide bridging GPR61 TM6 and TM7, with nearby motifs involved in activation switching highlighted.

Much like other active class A GPCR structures^18,20,21^, extracellular loop (ECL) 2 of GPR61 adopts a lid-like conformation over an orthosteric pocket, with its conformation stabilized by a conserved disulfide bond between Cys115^3.25^ (residue superscripts used throughout the manuscript indicate Ballesteros and Weinstein standardized GPCR residue position notations^22^, formatted with the TM helix number and associated standardized residue number separated by a period) and Cys195 of ECL2. Unlike many related receptors, however, the conformation of this loop largely occludes the orthosteric pocket, leaving a comparatively small and discontinuous pocket framed by TM1, 2, 3, and 7 and bisected by ECL2 (Fig. 1b). GPR61 lacks key ligand-interacting residues common to the orthosteric pockets of other biogenic amine receptors (e.g. Asp^3.32^, Asn^6.55^), though it retains conserved residue Phe304^6.52^. Diffuse density for the extracellular portion of TM1 suggests the pocket is plastic, and displacement of ECL2 by a ligand, as for instance in the case of rhodopsin^23,24^, could expose a larger and deeper pocket.

GPCR signaling through the cAMP pathway relies on binding of the C-terminal helix of Gα to an exposed intracellular pocket of the receptor formed primarily by activating movements of TM5 and TM6. This binding event triggers Gα to exchange GDP for GTP, leading to its activation. In the GPR61 structure, the C-terminal helix of Gαs binds within this pocket through a network of polar contacts involving residues of TM3, TM6, helix 8, and ICL2. On TM3, Tyr143^3.53^ and Val145^3.55^ form stabilizing hydrogen bonds with Gαs residues Gln390 and Gln384, respectively (Fig. 1c). Key to GPR61’s selectivity for Gαs are the interactions made by Gαs residue Glu392 with Arg346^8.48^ and Arg349^8.51^. Arg346^8.48^ makes a further interaction with the mainchain carbonyl of Gαs Arg389, forming a small hydrogen bonding network (Fig. 1d). Because Glu392 is found only in Gαs and G_olfα_, this hydrogen bonding pattern cannot be formed with other Gα proteins^25^. A number of non-polar interactions also drive interaction. A key residue of the D/ERY motif, Arg140^3.50^, forms an intricate ladder of hydrophobic interactions with Tyr143^3.53^ and Gαs residues Tyr391 and His387 (Fig. 1E). Though GPR61 lacks an ionic lock, Arg^3.50^ undergoes significant conformational rearrangement during activation of related receptors^26^, and this is also expected to be true of GPR61. Tyr228^5.58^ and Met231^5.61^ form further hydrophobic interactions with Leu393 of Gαs in the intracellular pocket formed by TM4, 5, and 6 (Fig. 1f).

The constitutive activity of GPR61 has been attributed to residues near its N terminus, with Val19 proposed to play a key role^9^. Some portion of this peptide might be expected to bind to the extracellular surface of the receptor, likely the orthosteric site, to accomplish activation. In our structure, the receptor’s initial 44 residues are unresolved, suggesting that any interactions of N-terminal residues with the receptor’s extracellular surface or orthosteric site are likely transient.

Efforts to capture this N-terminal region through crosslinking or addition of peptides derived from its sequence were unsuccessful. The bias of GPR61 toward constitutive activation does not appear to rely solely on its N-terminal peptide, however, as other key sequence and structural features are consistent with partial destabilization of the inactive state. In many class A receptors, Arg^3.50^ of the conserved D/ERY motif participates in a salt bridge, called the “ionic lock”, with an acidic residue (Asp or Glu) in position 6.30, which creates an energetic barrier to the movement of TM6 to accommodate binding of the Gα C terminus during activation^27^. In GPR61, Glu^6.30^ is substituted with a glycine (disordered in this structure), preventing the D/ERY motif (Glu139^3.49^, Arg140^3.50^, and Tyr141^3.51^) from making this interaction. Similarly, binding of Na^+^ ions to some class A GPCRs acts to stabilize the inactive unliganded state^28–30^, but GPR61 lacks key sodium-interacting residues (Ser^3.39^, Asn^7.45^, Ser^7.46^) and thus is not susceptible to this type of negative allosteric modulation. Interestingly, the defunct sodium binding site is juxtaposed with a disulfide bond between Cys299^6.47^ and Cys333^7.45^ that appears to be unique among GPCRs with known structures (Fig. 1g). A pair of cysteine residues in corresponding positions is found in only one other receptor, the class A P2Y purinoceptor 10, for which no structure is known, and AlphaFold predictions of both proteins fail to predict this disulfide. Other key features associated with activating conformational changes in GPCRs, including the Trp^6.48^ rotamer toggle switch and the N^7.49^PXXY motif^31^, are proximal to the disulfide, which may constrain the relative movements of TM6 and TM7 and thereby influence receptor modulation. Collectively, these features would be expected to weaken interactions known to stabilize the inactive conformation of related receptors, and thus may predispose GPR61 toward a more active basal conformation.

### Discovery of a potent and selective GPR61 inverse agonist

With the aim of identifying GPR61 inhibitors to treat cachexia, we initiated a high-throughput screening campaign leveraging Pfizer’s internal compound libraries against an assay measuring cAMP levels in a cell line overexpressing GPR61. Initial hits emerging from this screen were extensively optimized to yield a class of potent and selective sulfonamide-based GPR61 inverse agonists, represented here by compound 1, a tertiary sulfonamide (Fig. 2a). Consistent with its role in suppressing constitutive receptor activity, compound 1 increased the cell surface expression of a HiBit-tagged form of wild-type GPR61 (Fig. 2b, c). Compound 1 demonstrates excellent potency in the functional cAMP assay (IC_50_ = 10 nM) (Fig. 2d), with an inhibition profile consistent with that of an inverse agonist (Fig. 2e), and is selective for GPR61 among a panel of 9 GPCRs (off-target IC_50_ values >10 μM), with an excellent *in vitro* off-target profile (Extended Data Table 1). To better understand the molecular basis of Compound 1’s activity, we next pursued structural studies of GPR61 in its inactive state.

**Figure 2.**
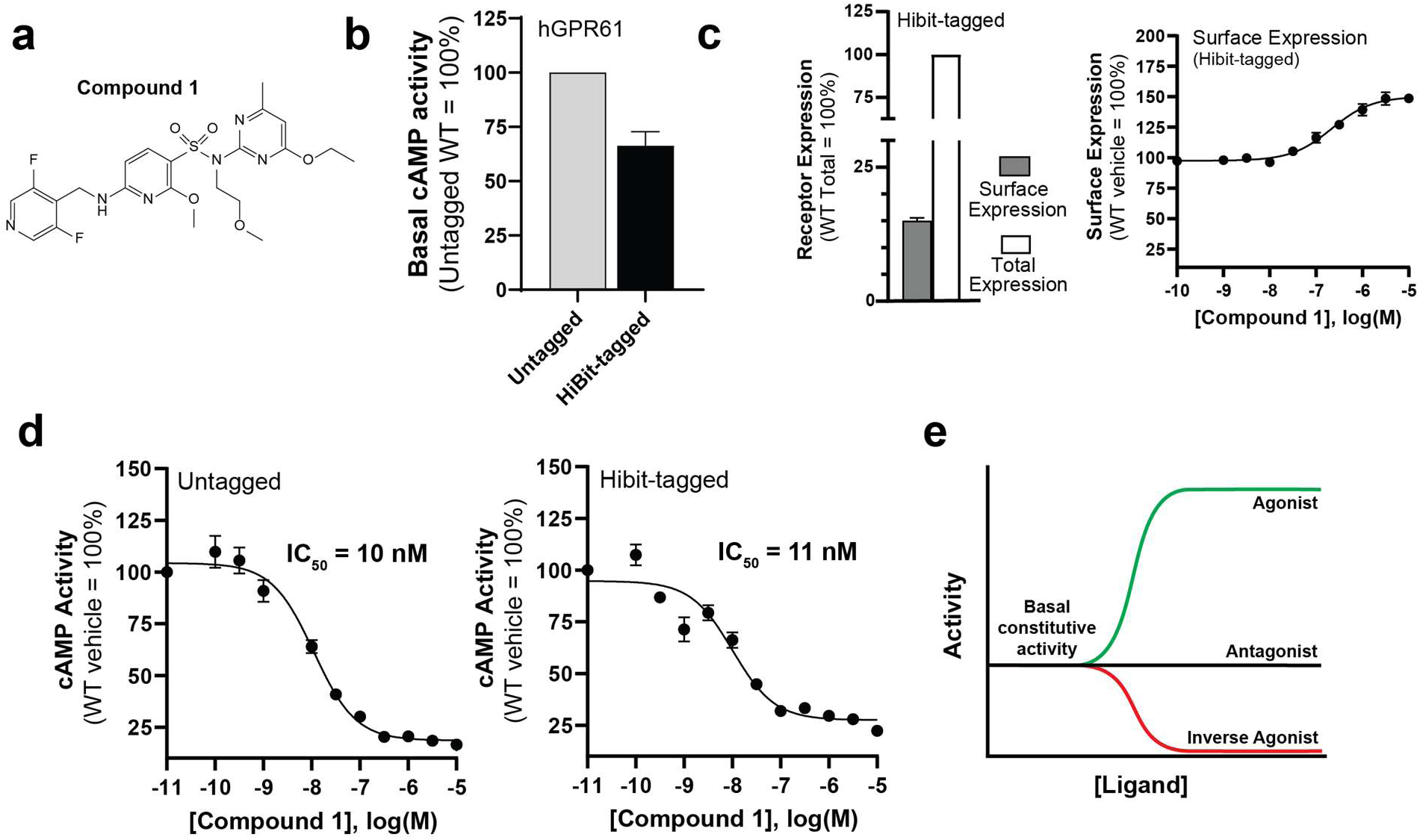
Inverse agonist compound structure and characterization. a. Structure of Compound 1. b. Relative activity in the cAMP assay of WT GPR61 and HiBit-tagged GPR61 used for measuring cell surface expression. c. Total and surface expression of GPR61 as measured using HiBit-tagged GPR61. d. Compound 1 cAMP assay inhibition curves and IC50 values for GPR61 (Untagged and HiBit-tagged). e. General schematic indicating the expected concentration-dependent responses to agonist, antagonist, and inverse agonist ligands. Data represent the mean ± S.E.M. from three independent experiments, each performed in duplicate.

### An AlphaFold-driven approach to inactive-state GPCR construct design

GPCR inverse agonists inhibit signaling by diverse mechanisms, acting at any of several allosteric sites on the receptor. Structural characterization is critical to understanding this process, so our next step was to pursue an inverse agonist-bound cryo-EM structure of GPR61. Structural characterization of inactive-state GPCRs by cryo-EM is made considerably more challenging by the loss of the heterotrimeric G protein, which acted as a fiducial for particle alignment in our active-state structure. To compensate for the loss of the G protein, we employed a strategy similar to that previously used to determine the structure of inactive Frizzled 5^32^, in which thermostabilized *E. coli* apocytochrome b562 RIL (BRIL^33^) was rigidly fused between TM5 and TM6, replacing intracellular loop 3 (ICL3). Because such dual helical fusions require careful optimization to ensure continuous helicity at both junctions, we combined knowledge gleaned from both our active-state GPR61 structure and the published Frizzled 5 structure to design a series of 25 GPR61-BRIL fusion constructs and screen their purified protein products by cryo-EM. A promising early construct yielded a model resolved to approximately 6 Å with clear separation of the seven TM helices, but efforts to improve the resolution of this construct further were unsuccessful. The subsequent availability of AlphaFold 2 allowed us to retrospectively compare the cryo-EM screening results for the fusion constructs against their corresponding AlphaFold predictions. We observed that constructs whose predictions showed rigid helical fusion points with high prediction confidence correlated with increased order in the results of our cryo-EM evaluation, building confidence in AlphaFold 2 as a tool for improved construct design. We employed this strategy to screen new construct designs *in silico* (Extended Data Fig. 1), selecting a subset of four for further characterization. AlphaFold predictions for all four indicated high-confidence helicity at the junctions between TM5/6 and the BRIL helices, but one construct, designated GPR61_IA_, stood out clearly as the best-ordered after screening by cryo-EM. This construct was used for all subsequent cryo-EM structural studies in combination with a previously described BRIL-binding Fab and a hinge-stabilizing nanobody^32,34^ as fiducials.

### Compound 1 binds a novel induced allosteric site in GPR61

The structure of apo-GPR61_IA_ was determined by cryo-EM to a nominal global resolution of 3.97 Å (Fig. 3a), but the region corresponding to the receptor was poorly resolved. This map was sufficient to identify and flexibly fit the TM helices to the density, but not to confidently place side chains. The positions of the TM helices revealed significant structural differences compared to the active-state receptor structure solved with the Gαs protein complex. The most pronounced of these was an inward rotation of the intracellular half of TM6 by about 12° toward TM1, 2, 3, and 7 relative to the active-state structure. This movement would cause TM6 to clash with the position of the C-terminal helix of GαS, leading to G protein complex dissociation and inactivation. While the structure’s resolution precluded a rigorous and detailed structural analysis, this shift in the conformation of TM6 is similar to that observed in other inactive-state receptors, and the receptor’s overall conformation is broadly consistent with the inactive conformation predicted for GPR61 in the GPCR database using AlphaFold 2-MultiState^35^.

**Figure 3.**
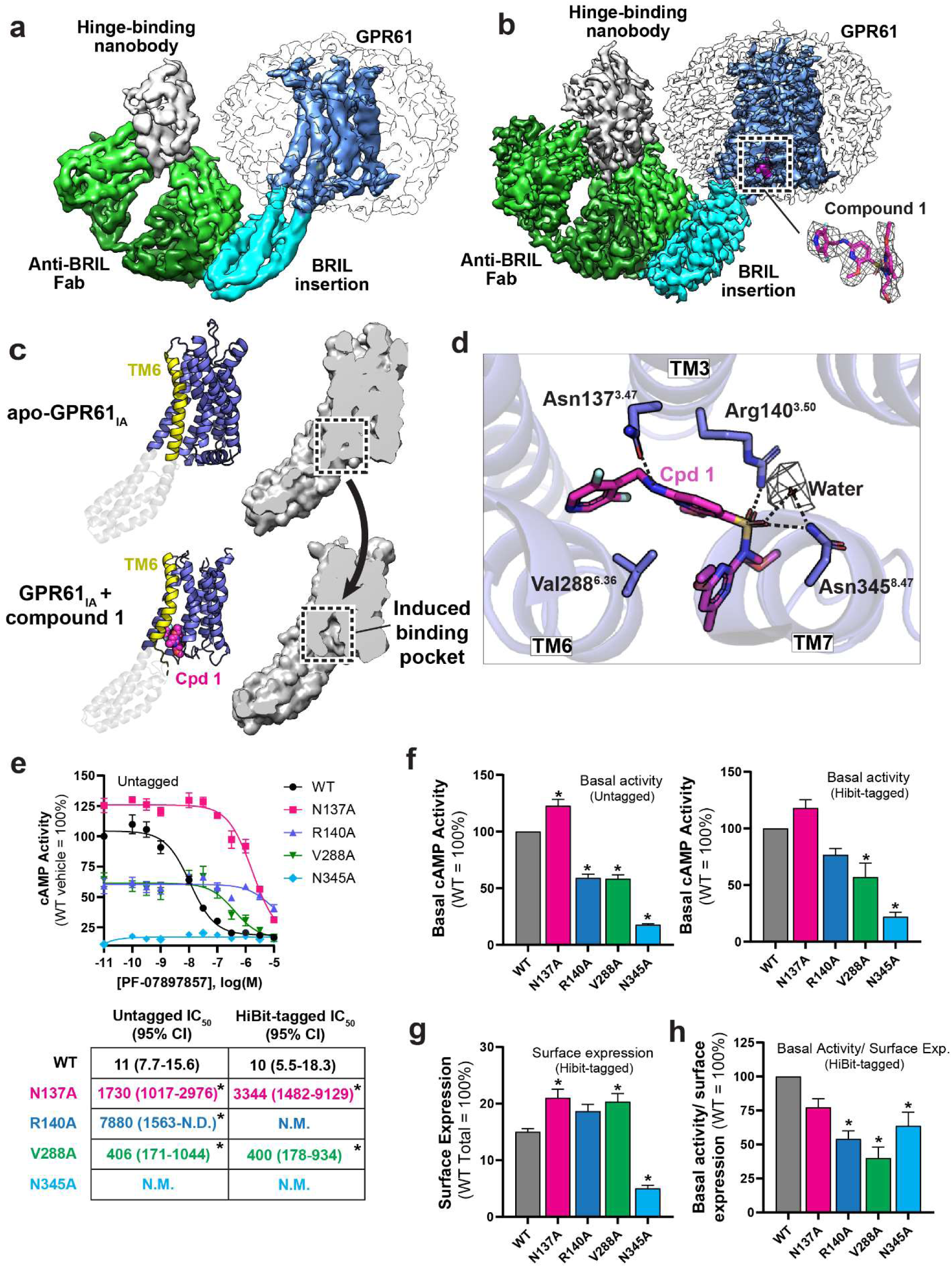
Structural and functional analysis of compound 1 binding to GPR61_IA_. a. Cryo-EM map of apo GPR61_IA_, colored by subunit. The sequence inserted into GPR61 (comprising BRIL and A2AR-derived linker sequences) is colored in cyan. b. Cryo-EM map of GPR61 _IA_ bound to compound 1, colored as in a. The compound 1 binding site is indicated by the dotted box. Inset shows compound 1, colored in magenta, fitted into its corresponding map density. c. A comparison of apo and compound 1-bound GPR61_IA_ conformations, showing the conformational changes induced by binding of compound 1. Ribbon diagrams, with TM6 highlighted, are shown at left, with a cutaway of the corresponding surface representation shown to the right. d. Compound 1 (Cpd 1) binding site, with key interaction residues. Map density for an ordered water is shown as dark gray mesh. Hydrogen bonds are indicated by dashed lines. e. GPR61 cAMP IC50 curves and values for WT GPR61 and the indicated mutants. f. Relative basal activity of untagged vs. HiBit-tagged GPR61 WT and mutants. g. Relative surface expression of GPR61 WT and mutants. h. Basal activity of GPR61 WT and mutants normalized to relative surface expression. Data represent the mean ± S.E.M. or 95% Cl from three independent experiments, each performed in duplicate. Statistical significance is indicated with an asterisk.

To obtain insights into the mechanism by which our inverse agonists inhibit GPR61 activity, we pursued a co-structure of GPR61_IA_ with compound 1 bound by cryo-EM. The presence of the inverse agonist significantly improved the order of the receptor relative to the apo structure, and the final map was resolved to 2.9 Å with excellent local resolution for the receptor (Fig. 3b). Modeling of the receptor into this map revealed a well-resolved region of unmodeled density whose shape is congruent to compound 1 (Fig. 3b, inset). Unexpectedly, compound 1 binds an induced allosteric pocket situated on the intracellular side of the receptor and flanked by TM helices 3, 5, 6, and 7. This site does not correspond to any other known GPCR ligand-binding site^36^, and it thus represents an entirely novel allosteric site for GPCR modulation. The formation of this induced binding pocket is enabled by a counter-intuitive conformational change in which the intracellular half of TM6 is forced outward relative to its position in the apo structure, nearly recapitulating the G protein-bound active form of the receptor (Fig. 3c). Alignment of this structure to the receptor’s active state gives an overall RMSD of 3.47 Å across the receptor, but an RMSD of only 0.44 Å for TM6. To accommodate this repositioning, the helical linkage between TM6 and BRIL is disrupted, with TM6 residues up to Lys284^6.32^ becoming disordered. This conformation is unique among inactive-state GPCR structures. While the artificial nature of the GPR61-BRIL fusion in GPR61_IA_ may give rise to concerns about whether the conformation of the apo protein is biologically relevant, the structural rearrangement induced by compound 1 suggests that the protein derived from this construct remains sufficiently flexible to accommodate ligand binding.

Key features of compound 1’s induced binding pocket reveal the basis of its potency. The bioactive conformation of compound 1 wraps around the side chain of Val288^6.36^, forming extensive stabilizing Van der Waals’ contacts (Fig. 3d). Its difluoropyridine group projects into a hydrophobic gap between TM5 and TM6, while the central linker’s methoxypyridyl is flanked by hydrophobic interactions with Val288^6.36^ and the β and γ carbons of Arg140^3.50^. The terminal methylpyrimidine projects toward the surrounding micelle by sandwiching between helices 6 and 7, while its ethoxy group extends toward Tyr341^7.53^ of the NPxxY motif (Extended Data Fig. 3).

Most critical for potency is compound 1’s sulfonamide moiety. Sulfonamides constitute a privileged chemotype among GPCR modulators, with many published examples^37–40^. The unique allosteric site bound by compound 1, however, defines a new class of sulfonamide GPCR inhibitors. The sulfonamide moiety of compound 1 forms key hydrogen bonds to Asn345^8.47^ and Arg140^3.50^, the key residue of the widely conserved D/ERY motif associated with activating conformational changes (Fig. 3d). Strong density for an ordered water is discernable in the map, coordinated between Asn345^8.47^ and the sulfonamide. Mutation of R140 or V288 to alanine made the receptor less sensitive to inverse agonism by compound 1 in the cAMP assay, while changing constitutive activity by less than 2-fold (Fig. 3e,f). In contrast, an N345A mutation significantly reduced the basal cAMP activity of the receptor, but additional investigation revealed this mutation to reduce the fraction of GPR61 at the plasma membrane (Fig. 3g). This may be attributable to the intracellular, solvent-exposed position of N345, whose mutation may impact receptor trafficking to the plasma membrane through the secretory pathway. When the N345A mutant’s basal activity was normalized to its cell surface expression, its activity was similar to that of the other mutants (Fig. 3h), but showed no sensitivity to compound 1 at up to 10 μM concentration.

The bioactive conformation of compound 1 likely also contributes to its potency. The strain energy of compound 1’s GPR61-bound conformation compared to the global minimum conformation is fairly small, estimated at ∼7.0 kcal/mol, with relatively small conformational differences. (Extended Data Fig. 2a,c). Torsional energy scans of the most disparate dihedral angles between the two conformations suggest very little strain associated with the adaptation of the difluoropyridyl tail to the binding pocket (Extended Data Table 2 and Extended Data Fig. 2a), but slightly larger strain energies are required for the amine and sulfonamide torsions that lead to the bound conformation (Extended Data Table 2 and Extended Data Fig. 2b,c). Since the overall strain energy is less than those of the individual torsion profile energy differences, the individual torsion scans likely overestimate the strain energy.

Compound 1 is selective for GPR61 (Extended Data Table 1), which may be attributed to the characteristics of the novel allosteric pocket. Compound 1 makes a key hydrogen bond through its secondary benzylic amine to the terminal amide oxygen of Asn137^3.47^ (Fig. 4a). The asparagine in this position is unique among known GPCRs, and in other receptors, substitutions in this position are non-conservative, with Ala and Ser being the most common replacements. As a key contributor to the compound’s potency, mutation of Asn137 would be expected to exact a large energetic penalty, reducing compound binding affinity considerably. Consistent with this hypothesis, mutation of Asn137^3.47^ to Ala in GPR61 reduced the potency of compound 1 about 100-fold relative to wild-type in the cAMP assay (Fig. 3e). Residues interacting with the sulfonamide moiety are key to the potency of these molecules are much more highly conserved and thus do not contribute significantly to selectivity. The remainder of the hydrophobic pocket is poorly conserved among other receptors, and sequence variation in these residues would be expected to alter the pocket’s shape complementarity to the compound for Van der Waals interactions, potentially reducing its affinity to varying degrees.

**Figure 4.**
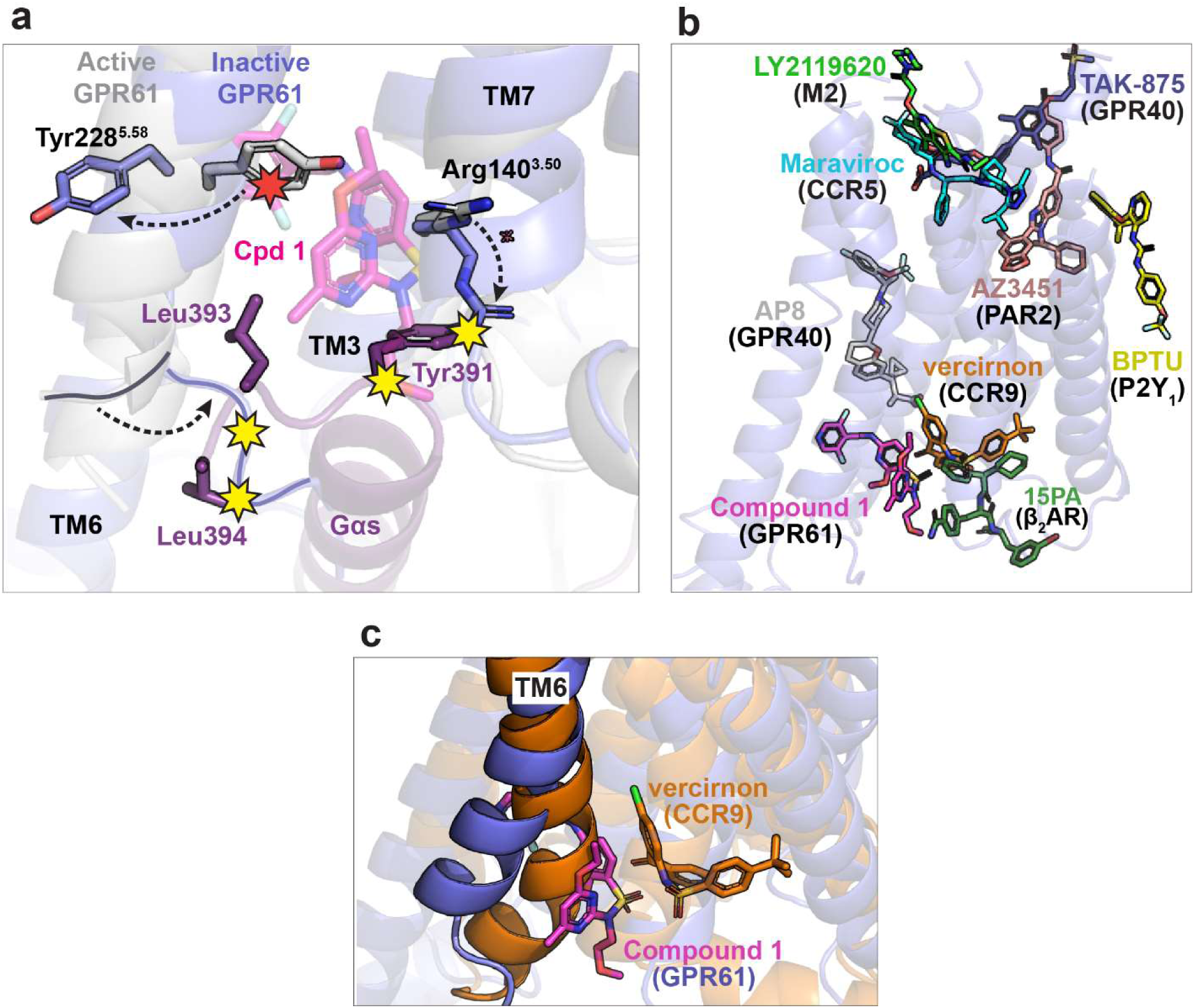
Analysis of compound 1 inverse agonist mechanism. a. Key residue clashes and conformational changes induced by binding of compound 1 (Cpd 1) to GPR61. The structure of active GPR61 (light grey) is overlaid with the compound 1-bound structure of inactive GPR61 (blue), with key residues highlighted in stick representation. Clashes with the compound are indicated by red stars, while clashes with Gαs induced by compound binding are indicated by yellow stars. b. Compound 1 defines a novel allosteric site and mechanism. The structure of compound 1-bound GPR61_IA_ is shown in ribbon representation, with published exemplars^44–51^ representing the known allosteric sites of class A GPCRs superimposed. c. Compound 1-bound GPR61_IA_ and vercirnon-bound CCR9^44^ structures, colored as indicated, are superimposed. Vercirnon occupies the known allosteric site nearest to that of compound 1. The different conformations of TM6 induced by these inverse agonists are highlighted.

### A novel inverse agonist mechanism

The induced binding pocket occupied by compound 1 gives rise to a new mechanism of GPCR inverse agonism. As discussed above, when bound to compound 1, the receptor adopts a conformation similar, but not identical, to that of the active receptor that nonetheless precludes the binding of Gαs necessary for activation of downstream signaling. Compound 1 acts as a “wedge”, binding in a pocket that partially overlaps that bound by Gαs in the active state. This wedge pushes TM6 outward compared to its position in the inactive-state apo structure, but not quite as far as seen in the active structure (Fig. 4a). This creates subtle differences in the positioning of the other flanking helices, which remodel the Gαs-binding pocket to reposition key hydrogen-bonding residues while the methoxyethyl substituent of the tertiary sulfonamide clashes with Tyr391 of Gαs.

To accommodate the terminal pyridine moiety of compound 1, the side chain of Tyr228^5.58^ on the displaced TM5 is flipped outward toward solvent, breaking a hydrophobic interaction with L393 of Gαs. Likewise, the position of TM6 differs slightly from the active-state structure, being positioned slightly closer to the neighboring helices and creating potential clashes with residues L393 and L394 of Gαs. L393 is framed by hydrophobic interactions with residues on TM5 and TM6, both of which are shifted by the presence of compound 1 (Fig. 4a). A significant outward shift of TM7 breaks interactions with Gαs residue Y391, while repositioning of Arg140^3.50^ disrupts the hydrophobic ladder of interactions with Gαs observed in the active-state structure and causes it to directly clash with Y391. Collectively, these structural changes remodel the Gαs-binding pocket to make G protein binding unfavorable. As a result, this inverse agonist mechanism paradoxically maintains a receptor conformation that more closely resembles the active state than the inactive state while still effectively blocking binding of the G protein.

Compared to the known GPCR allosteric sites discovered to date ^36,41^ (Fig. 4b), the site described here is novel. The most proximal known GPCR allosteric site is the highly conserved intracellular allosteric site observed for the β2AR, CCR, and CXCR receptors^42,43^, which neighbors, but does not overlap with that of compound 1. This site is flanked by helices 1, 2, 3, 6, and 7 and, like all other known allosteric inverse agonists, stabilizes the “inward” conformation of TM6 as its means of blocking Gα association, as exemplified by the structure of CCR9 bound to the inverse agonist vercirnon, another sulfonamide (PDB: 5LWE^44^) (Fig. 4c). Thus, compound 1 appears to be the first known example of a GPCR inverse agonist that induces a conformation more similar to active than inactive GPCRs.

## Discussion

GPR61 is an orphan class A GPCR with therapeutically relevant links to metabolic phenotypes. To date, the lack of structural information and reliable tool compounds for GPR61 have made comprehensive study of this receptor challenging. In the work described here, we introduce the first small-molecule inverse agonist, a sulfonamide with potent (10 nM) and selective activity against GPR61 (compound 1). Sulfonamides are a recurring chemotype among GPCR modulators, but compound 1 acts by an unprecedented mechanism, defining a new class of sulfonamide inverse agonists.

To better understand the basis of compound 1 activity, we used cryo-EM to determine the structure of GPR61 bound to compound 1 at 2.9 Å resolution, utilizing an efficient AlphaFold-driven in silico construct evaluation strategy to streamline the time-consuming process of experimental construct screening and optimization, providing significant savings in time and cost. revealing a mechanism completely unlike that of known GPCR inverse agonists (Fig. 5). Compound 1 binds to an unprecedented induced allosteric pocket that overlaps the binding site for Gαs and unexpectedly promotes a GPR61 conformation more similar to its active state than the inactive state. Compound 1 prevents Gαs binding and signal transduction both by sterically clashing with key residues of the Gαs C-terminal helix and by acting as a wedge to cause subtle conformational remodeling of the pocket, reducing its shape complementarity and eliminating favorable interactions with Gαs. This stands in contrast to all other known inverse agonists, which prevent Gαs binding by indirectly promoting a receptor’s inactive conformation. By directly blocking binding of Gαs to GPR61, compound 1 fits the mechanistic criteria for an inverse agonist, which blocks constitutive GPCR signaling, as opposed to an antagonist, which prevents receptor activation above its constitutive level.

**Figure 5.**
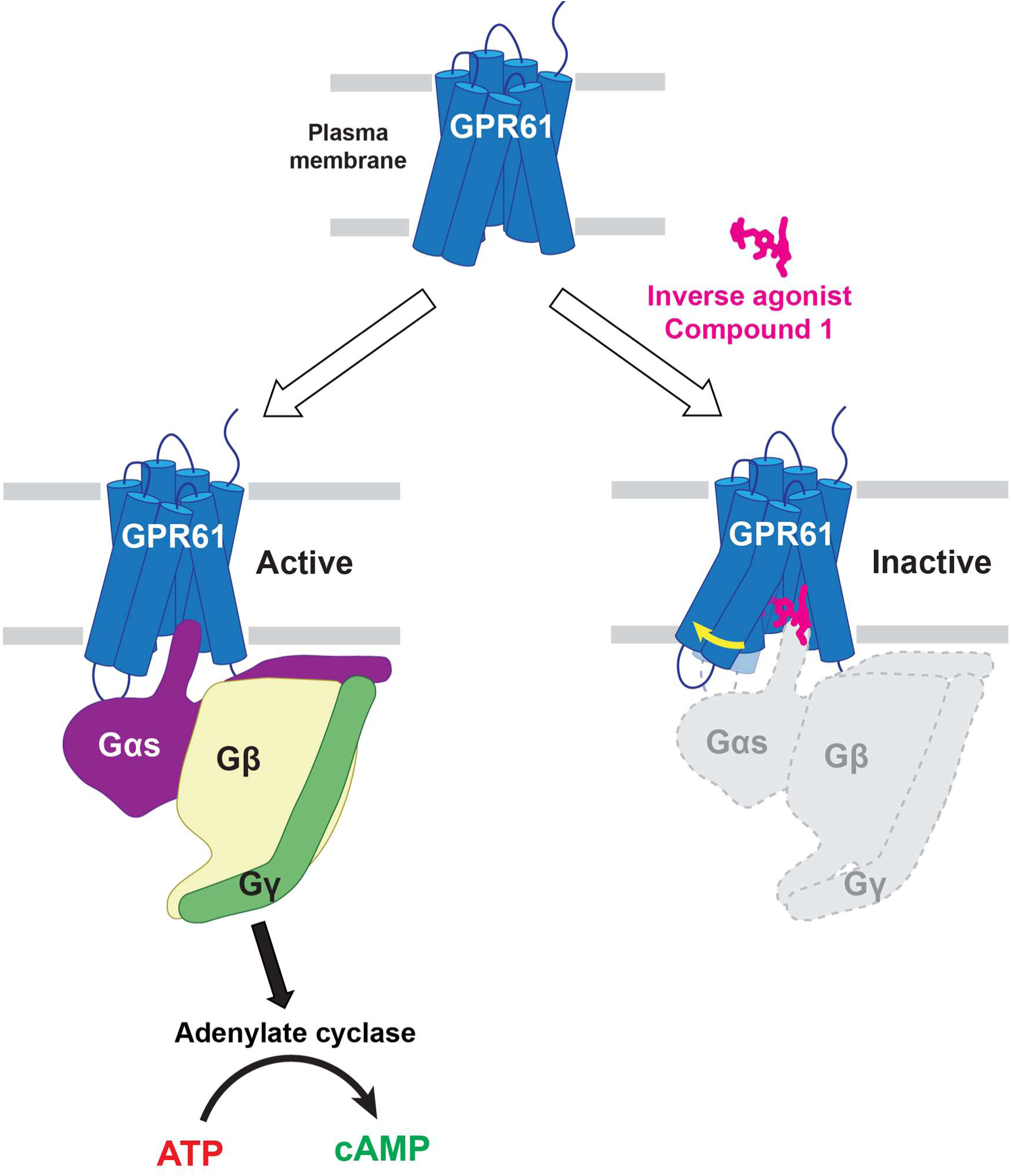
A new mechanism of allosteric GPCR inverse agonism. In its constitutively active state (top panel), GPR61 adopts a conformation that allows binding and nucleotide exchange of the G protein complex (bottom left panel) to stimulate cyclic AMP-mediated signaling through activation of adenylate cyclase. Inverse agonist compound 1 (magenta) binds to a novel intracellular allosteric pocket overlapping that bound by Gαs and acts as a wedge to remodel the flanking helices (yellow arrow), destroying the Gαs-binding pocket and creating direct clashes that prevent potential Gαs binding.

We have also determined unliganded structures of GPR61 in its active and inactive states, revealing the conformational changes associated with its activation, as well as the basis of its constitutive activity. In addition to providing mechanistic information, these structures provide a strategic platform for future mechanistic studies of GPR61 and modulators. The strategies we have used to enable these structures of GPR61, including a hybrid dominant negative-GαsiN18 chimera for the active state, and AlphaFold-informed construct design for the inactive state, will also be applicable to elucidating other GPCRs that still lack structural information. Together, the information we present here constitutes a toolbox for future study of GPR61 and other receptors, which will enable studies of receptor function and even facilitate receptor de-orphanization. Compound 1’s potency and excellent off-target profile make it a high-quality tool compound for future functional studies of GPR61 *in vivo* and provides a foothold, including a novel allosteric pocket, for future drug discovery efforts.

## Methods

### Construct design, protein expression and purification for cryo-EM study

The expression construct for human GPR61 in the active conformation (termed hGPR61-dnGαs/iN18), was designed with an HA signal peptide, FLAG tag, TEV protease cleavage site, BRIL (cytochrome b562 RIL), and PreScission protease cleavage site at the GPR61 N terminus, and with the GPR61 C terminus fused via a (GSS)_9_ linker to a novel chimeric dominant negative Gαs/iN18 (combining truncation Gαi N18, Gαs N26^14^ with a previously described dominant negative version of Gαs^13^). The expression construct for human GPR61 inactive conformation (termed hGPR61_ICL3_BRIL) was designed with an HA signal peptide at the N-terminus and a C-terminal FLAG tag, a modified BRIL sequence was inserted between ICL3 residues R233 and K285.

GPR61 constructs were expressed in *Spodoptera frugiperda* (Sf9) insect cells using the Bac-to-Bac system. Briefly, the receptor was expressed for 48-72 h after infection and harvested cells were stored at -80°C until further use.

For purification of GPR61 constructs, membranes were isolated by lysing the cells and ultra-centrifuging lysates for 45 min. The membranes were resuspended and washed in 500 mM NaCl, 50 mM HEPES pH 7.5, and EDTA-free cOmplete Protease Inhibitor (Roche), and ultra-centrifuged as before. Membranes were solubilized with 1% LMNG (Anatrace) and 0.2% CHS (Anatrace) in 500 mM NaCl and 50 mM HEPES pH 7.5, 100 μM TCEP, and EDTA-free cOmplete Protease Inhibitor (Roche). The solubilized protein was isolated using FLAG (Sigma) affinity gel, then further purified by size exclusion chromatography on a Superose 6 (Cytiva) column equilibrated in a buffer containing 0.001% LMNG, 0.0002% CHS, 150 mM NaCl, 25 mM HEPES pH 7.5, 100 μM TCEP. The fractions were analyzed by SDS-PAGE and fractions corresponding to monomeric GPR61 were pooled.

N-terminally 6xHis-tagged human Gβ1 (S2-N340) and human Gγ2 (M1-C68) were synthesized and separately cloned into pFastBac1 (Thermo Fisher) and expressed as described above. Cells were lysed in buffer containing 20 mM HEPES pH 8, 150 mM NaCl, 1 mM TCEP, 0.5 mM EDTA, EDTA-free protease inhibitor cocktail (Roche), and benzonase, then isolated using a HisTrap Crude FF column (GE Healthcare) and eluted with 200 mM imidazole in the same buffer. Fractions containing the heterodimer were pooled and dialyzed overnight against 20 mM HEPES pH 8, 1 mM TCEP, 0.5 mM EDTA. The protein was bound to a HiTrap Q column (Amersham Biosciences) equilibrated in the same buffer and eluted with a linear NaCl gradient. Fractions containing the heterodimer were pooled and subjected to gel filtration on a Superdex 75 column in buffer containing 20 mM HEPES pH 8, 150 mM NaCl, 1 mM TCEP, 1 mM EDTA. Peak fractions were pooled, snap-frozen, and stored at -80°C until use.

To prepare the hGPR61-dnGαs/iN18-G protein complex, the purified hGPR61-dNGαs/iN18 fusion protein was incubated with an excess of Gβ, Gγ, scFv16 and apyrase (New England Biolabs), for 1 h on ice following published protocols ^14,15^. The complex was used directly for cryo-EM grid preparation without concentration.

For the preparation of the inverse agonist complex, the purified hGPR61_ICL3_BRIL fusion was incubated on ice for one hour with an excess of anti-BRIL Fab and anti-Fab nanobody according to published protocols^32^. The complex was concentrated and purified by size exclusion chromatography using a Superose 6 Increase 5/150 GL (Cytiva) in a buffer containing 0.001% LMNG, 0.0002% CHS, 150 mM NaCl, 25 mM HEPES pH 7.5, 100 μM TCEP. The fractions corresponding to the ternary GPR61+Fab+Nb complex were pooled. The complex was incubated with 100 μM of inverse agonist compound 1 overnight and was used directly without further concentration steps for cryo-EM grid preparation.

### Cryo-EM Sample Preparation

Purified protein samples were subjected to centrifugation at 13,200xg for 10 minutes to remove aggregates. Gold Quantifoil R1.2/1.3 200 mesh grids were made hydrophilic by glow discharge in residual air at 15 mA for 30 seconds using a Pelco Easiglow. In a Vitrobot Mark IV operated at 4°C and 100% humidity, 4 μl of sample supernatant was applied to a grid, then blotted away from both sides before being vitrified by plunge-freezing in liquid ethane cooled by liquid nitrogen. Vitrified grids were stored under liquid nitrogen until imaging.

### Cryo-EM Data Collection and Processing

Grids were imaged in a Titan Krios G2 transmission electron microscope operated at 300 kV equipped with a Falcon 4i direct electron detector and Selectris X imaging filter. All screening and data collection were performed in EPU (Thermo Fisher Scientific). Movies in EER format were collected at 215,000x magnification (0.59 Å magnified pixel size at the specimen level) with a total electron dose of 50 e^-^/Å^2^.

For the hGPR61-dnGαsiN18/Gβ/Gγ/scFv16 complex, a dataset of 20,635 movies was collected. Movies were subjected to patch motion correction and patch CTF correction in CryoSPARC 3.3.1 ^52^, followed by blob-based autopicking. 2,768,397 particles were extracted then subjected to 2D classification. 2D classes showing signs of secondary structure (255,448 particles) were subjected to a second round of 2D classification into 200 classes. 188,276 particles were subjected to 3D ab initio modeling in 4 classes. The best model, comprising 52,887 particles, was subjected to non-uniform 3D gold-standard refinement and reached a final resolution of 3.47 Å, based on the FSC=0.143 criterion.

For GPR61-BRIL fusion constructs, screening datasets of 5,000 movies were initially collected and processed in CryoSPARC as described below up to 2D classification. The appearance and quality of 2D classes were used to compare and evaluate constructs. Once the final construct was selected, a dataset of 16,126 movies was collected in the presence of inverse agonist compound 1, while a dataset of 10,000 movies was collected from the equivalent sample without compound to solve the apo structure. Movies were subjected to patch motion correction and patch CTF correction in CryoSPARC 3.3.1, followed by blob-based autopicking. Particles were extracted and subjected to 2D classification into 200 classes. 2D classes resembling a micelle with protruding density were manually selected and subjected to a second round of 2D classification in 200 classes. Well-defined 2D classes representing all discernable particle views were then used for template-based picking against the dataset. Picked particles were extracted as before and subjected to two rounds of 2D classification into 200 classes. Particles from well-resolved 2D classes were fed into 3D ab initio modeling followed by non-uniform gold-standard 3D refinement.

### Model building and refinement

For each of the GPR61 structures reported here, an atomic model predicted by AlphaFold 2^53,54^ (wild-type GPR61 for the active state structure and hGPR61_ICL3_BRIL fusion construct sequence for the inactive state) was rigid-body fitted into the map density. The model was successively hand-built into the map using Coot (v0.9.8.1^55^) in alternation with real-space refinement in Phenix to produce the final model. Starting models of Fab24 BAK5 and the hinge-binding nanobody were derived from PDB entry 6WW2^32^, while initial models of dnGαs/iN18, Gβ, Gγ, and scFv16 were created by modifications of PDB entry 3SN6 ^18^. The full cryo-EM data processing workflow and validation metrics (Extended Data Fig. 4, 5, 6) and the model refinement statistics (Extended Data Table 3) can be found in the supplementary materials. Figures based on the structure were produced in PyMol version 2.5.4, UCSF Chimera version 1.16^56^, and ChimeraX version 1.4^57^.

### Computational Chemistry

The strain energy of compound 1 was computed by taking the energy difference between the cryo-EM (local minimum) and global minimum conformations.

The energy of the local minimum conformation was determined by minimizing the cryo-EM structure of compound 1 using a 10 kJ/mol Å^2^ constraint on all torsions. The energy of the global minimum conformation was determined by performing a conformational search of compound 1 and selecting the lowest energy conformation. Torsional energy scans were performed by defining the dihedral angle(s) of interest in compound 1 and increasing them from 0° to 360° by increments of 10° using Coordinate Scan. All calculations were run using Macromodel^58^ in Schrödinger 2021-2^59^ using a dielectric constant to approximate water using default options unless otherwise specified. All conformational energies were determined using the OPLS4 force field,^60^ which was customized using the Force Field Builder panel for missing ligand torsion parameters.

## Author Contributions

J.A.L. prepared cryo-EM grids, collected and processed cryo-EM data, and performed model building. J.A.L., J.M.D, and S.H. analyzed the structures. J.A.L., F.R., E.C., and J.M.D. ran AlphaFold 2 predictions and designed constructs. F.R. and E.C. produced constructs, generated baculoviruses, and expressed constructs for cryo-EM. J.M.D. optimized protein purification and purified proteins for cryo-EM studies. J.M.D., F.R., J.A.L., A.E.V., A.M., and E.C. purified proteins for inactive-state GPR61 construct screening. R.O. performed and analyzed cAMP activity assays. J.P.F., J.X.K, and E.H. generated constructs and cell lines for pharmacological studies and performed and analyzed receptor expression studies. Y.Z., J.T., G.L., B.K., R.U., and L.Z. contributed to design efforts leading to compound 1. B.K. performed compound 1 energy calculations. A.M.D.S., E.F., and D.Z. enabled synthesis of compound 1. Y.Z. supervised the project chemistry. S.H. supervised efforts related to cryo-EM. J.A.L., J.M.D., and S.H. wrote the manuscript, with analysis and input from all authors.

## Acknowledgments

The authors gratefully acknowledge K. Sumner, S. Rotstein, D. Tse, and J. Yee for ongoing support of our cryo-EM computing infrastructure.

