## supplementary for "An inverse agonist of orphan receptor GPR61 reveals a novel allosteric mechanism"

**Extended Data Figure/Table Legends**

**Extended Data Table 1: Compound 1 Selectivity.** Stimulatory and/or inhibitory activity of compound 1 against GPCRs and a panel of drug safety target assays.

**Extended Data Table 2: Conformational Energetics of Compound 1.** Select dihedral angle differences between the global minimum conformation of compound 1 and the bound cryo-EM conformation with estimated energy differences based on the OPLS4 force field.

**Extended Data Table 3: Cryo-EM Data Collection, Processing, and Refinement Statistics.**

**Extended Data Figure 1: AlphaFold-guided GPR61-BRIL Construct Design.** a. General schematic for designing constructs. BRIL was fused, with or without linker sequences, to replace intracellular loop 3 of GPR61 for continouous helical extensions to GPR61 TM5 and TM6. b. General workflow for AlphaFold-based screening of constructs. AlphaFold predictions were used to bin construct designs based on the quality of helical fusions to TM5 and TM6 of GPR61. Predictions with relatively straight, helical fusions for both helices were selected for screening by cryo-EM. Cryo-EM screening enabled identification of a construct suitable for scale-up and full 3D reconstruction.

**Extended Data Figure 2: Compound 1 Energy Calculations.** Energy profile of the difluoropyridyl tail dihedral angle 1. Dihedral angles of the global minimum conformation (236.0°) and the cryo-EM conformation (255.3°) are indicated by the solid and dashed lines, respectively. **B.** 2D energy profile of the amine dihedral angles 2 and 3. The magenta circle indicates the global minimum conformation (74.9° and 0.3°, respectively) and the gray star indicates the cryo-EM conformation (122.1° and 330.2°, respectively). **C.** Energy profile of the sulfonamide dihedral angle 4. Dihedral angles of the global minimum conformation (125.9°) and the cryo-EM conformation (106.7°) are indicated by the solid and dashed lines, respectively.

**Extended Data Figure 3: Compound 1 Protein-Ligand Interactions.** Residues proximal to compound 1 in the cryo-EM structure are indicated by labels. Green arrows indicate hydrogen bonds, with the arrow pointing from donor to acceptor.

**Extended Data Figure 4: Cryo-EM Data Processing Workflow for Active-State GPR61-dnGαs/i chimera-Gβγ.** a. Data processing workflow. b. Fourier shell correlation curves from gold-standard refinement. c. Samples of map density with fitted model segments.

**Extended Data Figure 5: Cryo-EM Data Processing Workflow for Apo-GPR61$_{IA}$.** a. Data processing workflow. b. Fourier shell correlation curves from gold-standard refinement. c. Samples of map density with fitted model segments.

**Extended Data Figure 6: Cryo-EM Data Processing Workflow for GPR61$_{IA}$ + Compound 1.** a. Data processing workflow. b. Fourier shell correlation curves from gold-standard refinement. c. Samples of map density with fitted model segments.

#### Extended Data Table 1

| Compound 1 GPCR Selectivity | | |
|---|---|---|
| GPCR Target | Agonist EC50 (nM) | Antagonist IC50 (nM) |
| Alpha adrenergic receptor 1a | >10,000 | >10,000 |
| Beta-2 adrenergic receptor | >10,000 | >10,000 |
| Cannabinoid receptor 1 | >10,000 | >10,000 |
| Dopamine receptor 1 | >10,000 | >10,000 |
| Histamine receptor 1 | >10,000 | >10,000 |
| Mu opioid receptor | >10,000 | >10,000 |
| Serotonin receptor 2b | >10,000 | >10,000 |
| Muscarinic receptor 1 | >10,000 | >10,000 |
| Muscarinic receptor 3 | >10,000 | >10,000 |

| Transporter Targets | IC50 (nM) |
|---|---|
| Serotonin Transporter | >10,000 |
| Norepinephrine Transporter | >10,000 |
| Dopamine Transporter | >10,000 |

| Phosphodiesterase Targets | IC50 (nM) |
|---|---|
| Human PDE1B1 | >40000 |
| Human PDE2A1 | >200000 |
| Human PDE3A1 | 18357 |
| Human PDE4D3 | >35018 |
| Human PDE5A1 | 154918 |
| Bovine PDE6 | >200000 |
| Human PDE7B | 6273 |
| Human PDE8B | 9071 |
| Human PED9A1 | >200000 |
| Human PDE10A1 | >40000 |
| Human PDE11A4 | >40000 |

| Epigenetic Targets | IC50 (nM) |
|---|---|
| BRD4 (Binding) | >25000 |

| Enzyme Targets | IC50 (nM) |
|---|---|
| Acetylcholinesterase | >50000 |

| Ion Channel Targets | Patch IC50 (nM) | Binding Ki (nM) |
|---|---|---|
| hERG | 52904 | >80000 |

| Ion Channel Targets | %Activation@10μM | %Inhibition@10μM | EC50 (nM) | PAM EC50 (nM) |
|---|---|---|---|---|
| GABAa | 0 | 4 | >100000 | >100000 |

| Ion Channel Targets | IC50 (nM) |
|---|---|
| Calcium Channel Cav1 | 44787 |
| Sodium Channel Nav1 | >100000 |
| Sodium Channel Nav1.5 (Peak) | >100000 |

| | Period | | |
|---|---|---|---|
| Ion Channel Extended Protocol IC50 (nM) | 1st | 2nd | 3rd |
| hERG | 36519 | 28050 | 18705 |
| Cav1.2 | 21061 | 10729 | 5926 |
| Nav1.5 (Peak) | >100000 | >100000 | >100000 |

**Extended Data Table 2**

| Dihedral Angle Name | Cryo-EM Dihedral Angle (°) | Global Minimum Dihedral Angle (°) | ΔG (kcal/mol) |
|---|---|---|---|
| Difluoropyridyl tail torsion **1** (NH-C-C-CF) | 255.2 | 236.0 | 0.6 |
| Amine torsion **2** | 122.1 | 74.9 | 4.9 |
| Amine torsion **3** | 330.2 | 0.3 | |
| Sulfonamide torsion **4** | 106.7 | 125.9 | 2.2 |

#### Extended Data Table 3. Cryo-EM data collection, refinement and validation statistics

| | GPR61-dnGαs/i chimera-Gβγ<br>(EMDB-xxxx)<br>(PDB xxxx) | GPR61-BRIL+ Compound 1<br>(EMDB-xxxx)<br>(PDB xxxx) | GPR61-BRIL apo<br>(EMDB-xxxx)<br>(PDB xxxx) |
|---|---|---|---|
| **Data collection and processing** | | | |
| Magnification | 215,000x | 215,000x | 215,000x |
| Voltage (kV) | 300 | 300 | 300 |
| Electron exposure (e–/Å$^2$) | 50 | 50 | 50 |
| Defocus range (μm) | -2.4 to -0.6 | -2.4 to -0.6 | -2.4 to -0.6 |
| Pixel size (Å) | 0.59 | 0.59 | 0.59 |
| Symmetry imposed | C1 | C1 | C1 |
| Initial particle images (no.) | 2,768,397 | 4,736,846 | 2,955,528 |
| Final particle images (no.) | 52,887 | 170,451 | 92,842 |
| Map resolution (Å)<br>FSC threshold | 3.47<br>(0.143) | 2.90<br>(0.143) | 3.97<br>(0.143) |
| Map resolution range (Å) | | 5.85-2.60 | |
| **Refinement** | | | |
| Initial model used (PDB code) | AF_AFQ9BZJ8F1,<br>3SN6 | AlphaFold,<br>6WW2 | AlphaFold,<br>6WW2 |
| Model resolution (Å)<br>FSC threshold | 3.5<br>0.5 | 3.1<br>0.5 | 4.0<br>0.5 |
| Model resolution range (Å) | | | |
| Map sharpening *B* factor (Å$^2$) | 82.7 | 100.5 | 103.1 |
| Model composition<br>Non-hydrogen atoms<br>Protein residues<br>Ligands | <br>8,467<br>1,090<br>0 | <br>6,809<br>882<br>1 | <br>6,304<br>399<br>0 |
| *B* factors (Å$^2$)<br>Protein<br>Ligand | <br>66.02<br>N/A | <br>37.80<br>51.53 | <br>234.12<br>N/A |
| R.m.s. deviations<br>Bond lengths (Å)<br>Bond angles (°) | <br>0.008<br>1.084 | <br>0.03<br>0.604 | <br>0.034<br>1.147 |
| Validation<br>MolProbity score<br>Clashscore<br>Poor rotamers (%) | <br>2.22<br>16.61<br>0 | <br>1.72<br>6.15<br>0 | <br>2.28<br>23.32<br>0 |
| Ramachandran plot<br>Favored (%)<br>Allowed (%)<br>Disallowed (%) | <br>91.56<br>8.07<br>0.38 | <br>94.33<br>5.44<br>0.23 | <br>93.70<br>6.05<br>0.25 |

#### Extended Data Figure 1

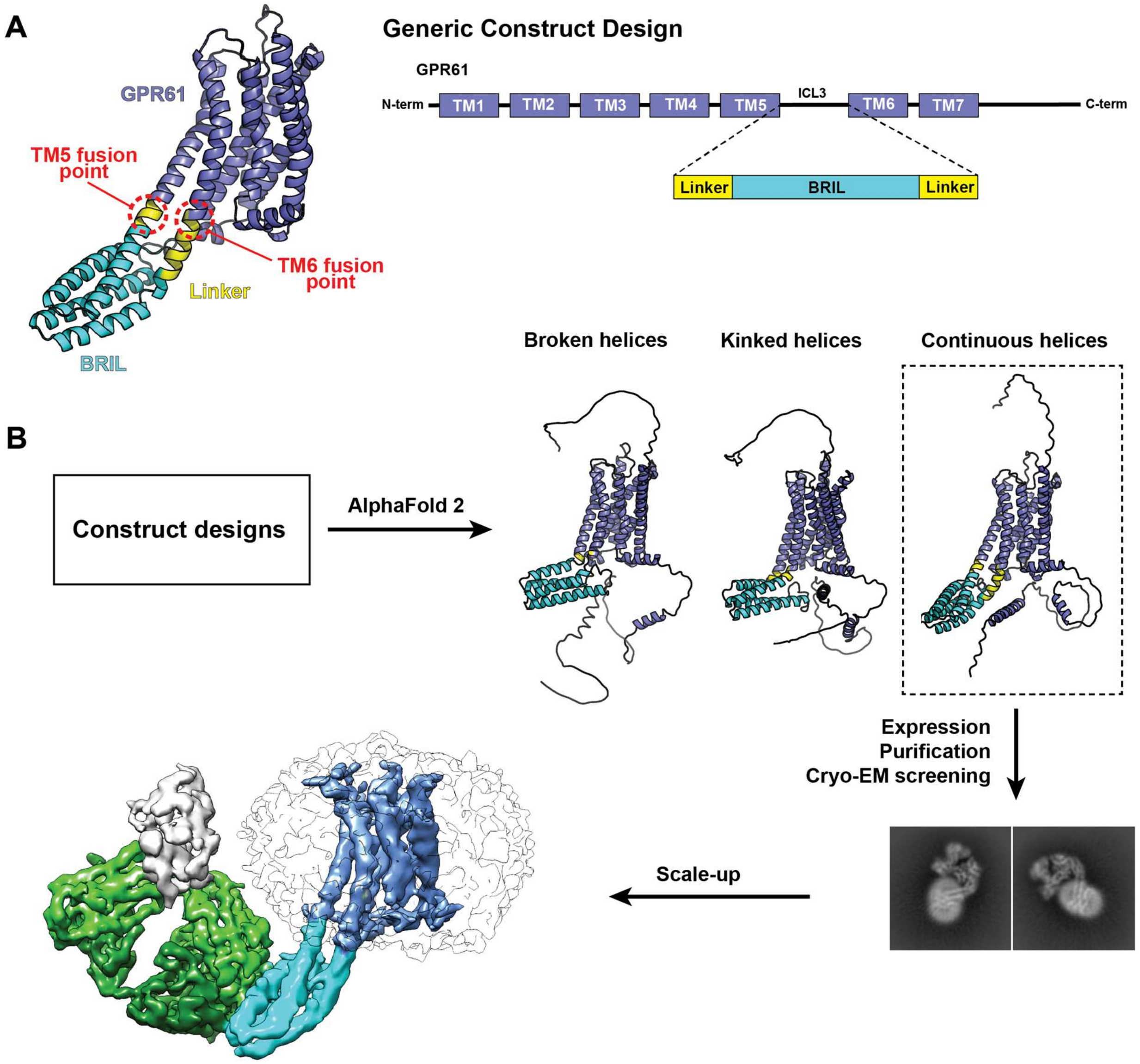

46

#### Extended Data Figure 2

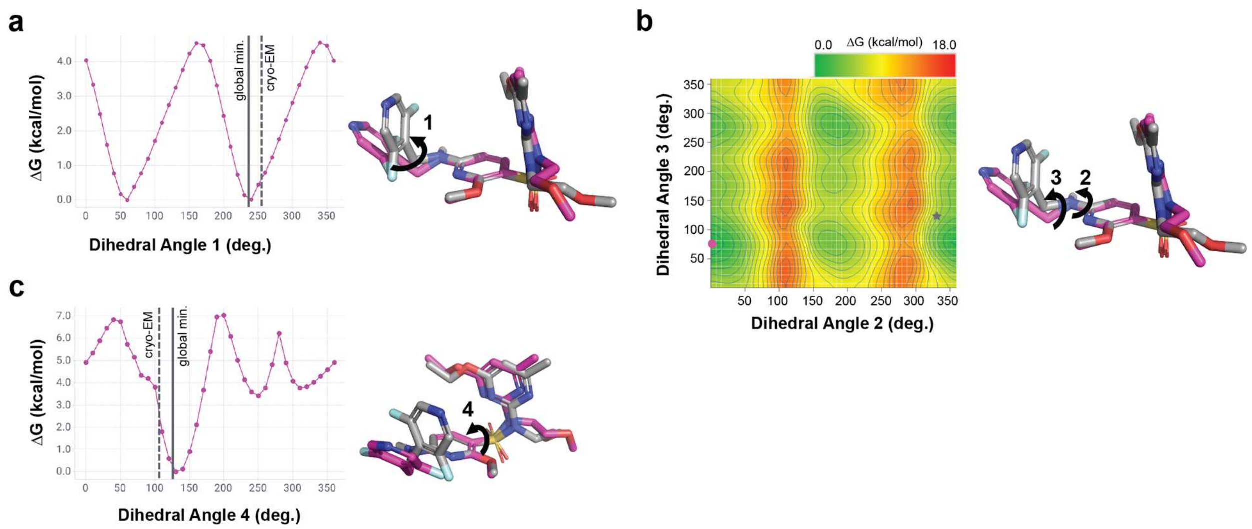

### Extended Data Figure 3

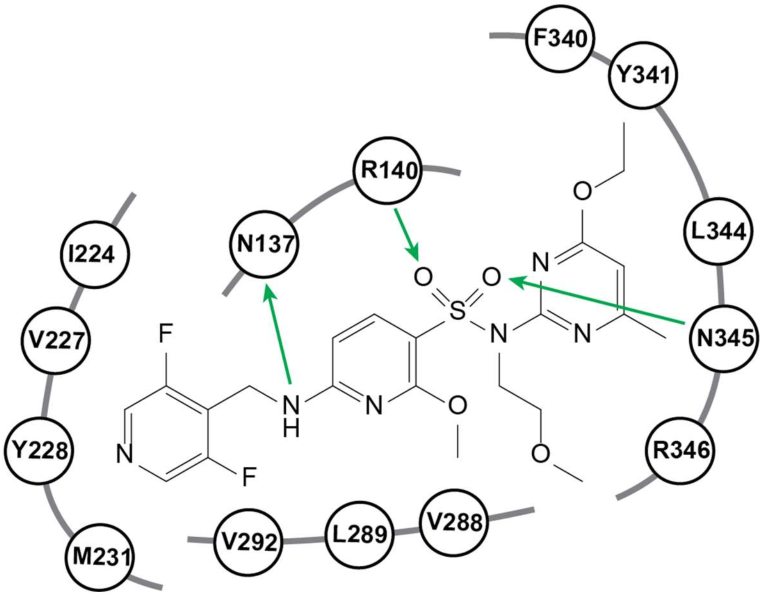

#### Extended Data Figure 4

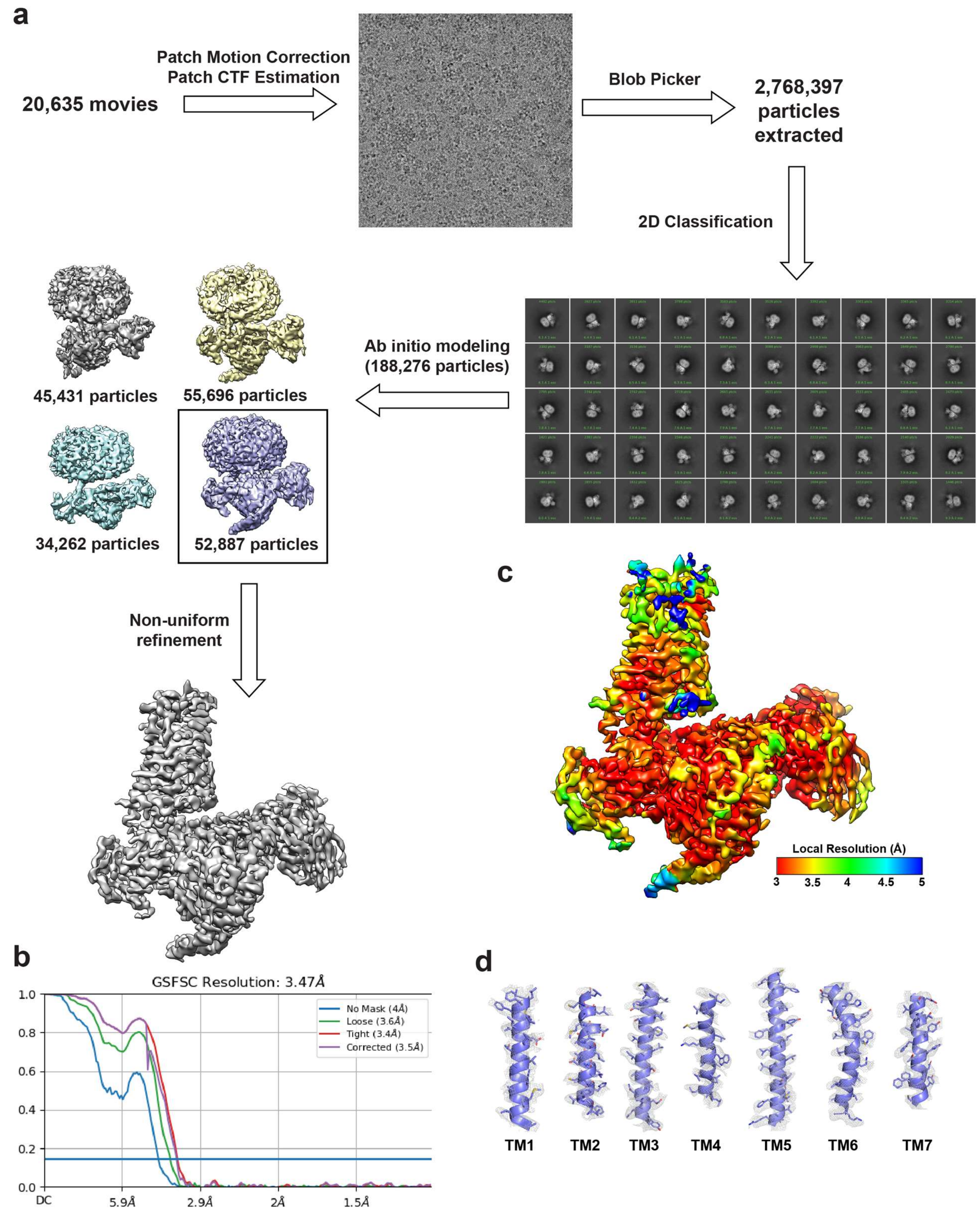

#### Extended Data Figure 5

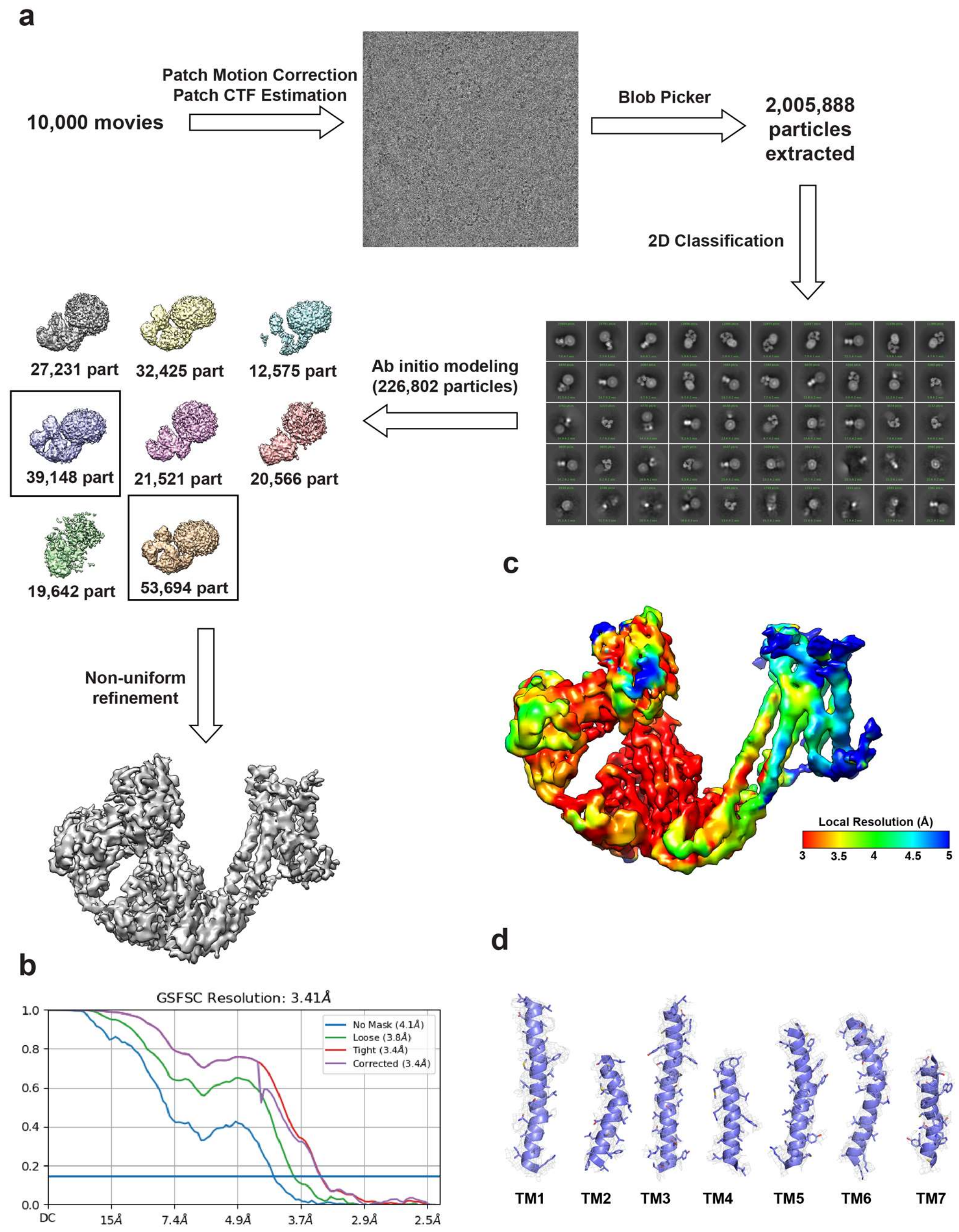

Lorem ipsum

#### Extended Data Figure 6

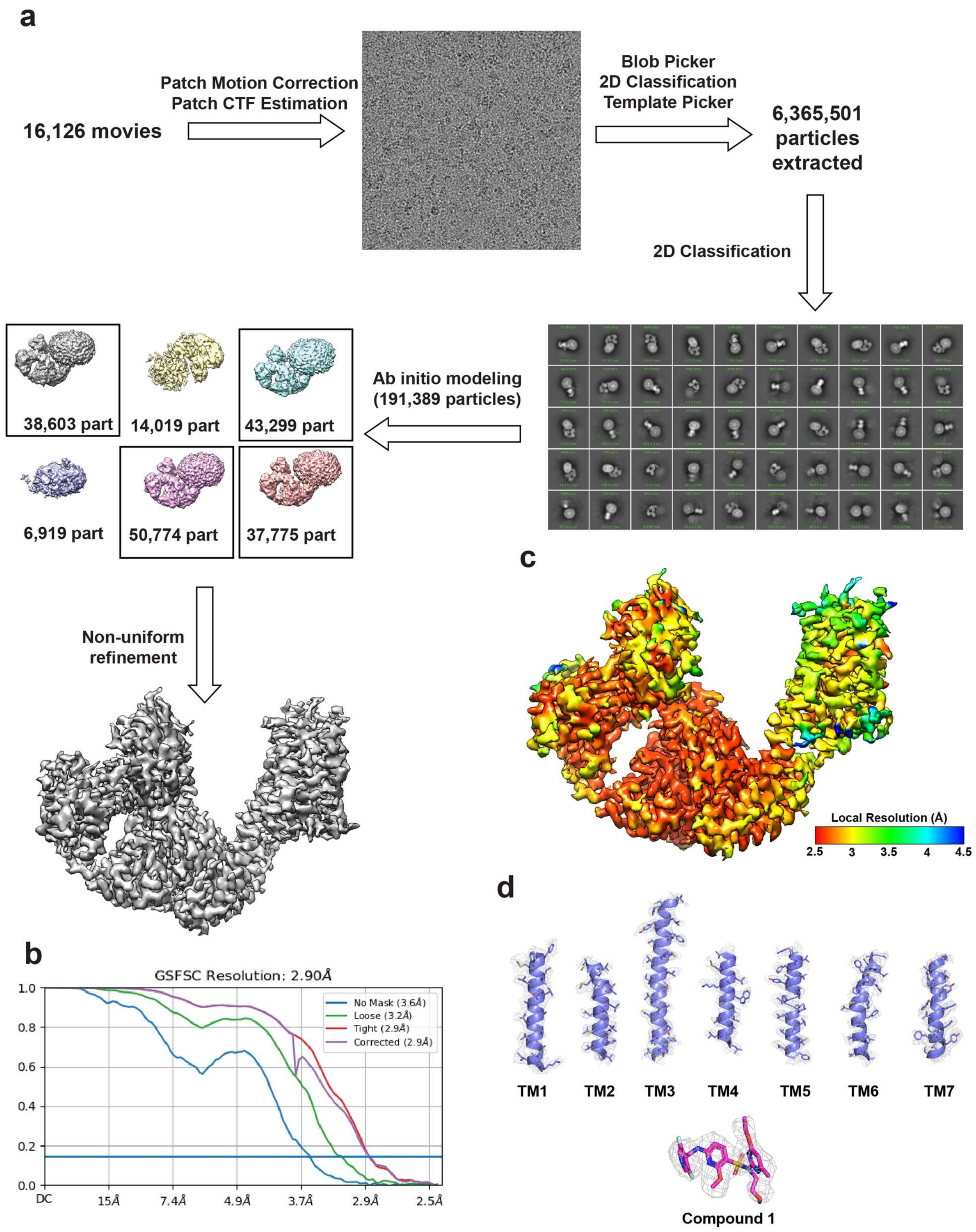

51

#### Extended Data Methods

##### Compound 1 Selectivity Analyses:

##### GPCR selectivity panel:

Adrenergic alpha1a, dopamine 1, histamine 1, muscarinic 1, muscarinic 3 and serotonin 2b receptor activities were measured using FLIPR® Calcium Assays.

Cells used in the assay were stably transfected with the receptor of interest (adrenergic alpha1a, dopamine 1, histamine 1, muscarinic 1, muscarinic 3 and serotonin 2b). Activation of the receptor by an agonist in this assay system results in an increase in intracellular calcium levels which is measured using a calcium specific dye. Cells were plated at 7,500 cells per well (50 µL per well) in black walled clear bottomed 384-well plates 24 h prior to running the assay. Medium was removed from the plates and 80 µL of Hanks balanced salt solution (HBSS)/HEPES containing Calcium 5 dye (Molecular Devices, Sunnyvale CA, USA; Cat # R8186) and probenecid (1.25 mM) was added to each well and the plate was returned to the incubator for 1 h to allow dye loading. Compound solution (10 µL) was added to each well by the FLIPR Tetra® instrument (Molecular Devices, Sunnyvale CA, USA) to measure agonist activity of the compound by measuring the change in fluorescence from baseline over a 60 second period (Excitation 470-495 nm; Emission 515-575 nm). Subsequently 10 µL of agonist ($EC_{80}$ value) was added to each well by the FLIPR Tetra® instrument to evaluate antagonist activity, with the change in fluorescence from baseline being measured over a 60 second period.

Adrenergic beta 2, cannabinoid 1 and mu opioid receptor activities were measured using Beta-Arrestin Assays.

The beta-arrestin assay relies on enzyme fragment complementation with the respective stably transfected GPCR (adrenergic beta 2, cannabinoid 1 and mu opioid) being tagged with an inactive portion of the enzyme β-galactosidase and a co-transfected β-arrestin that is tagged with the complementary portion of β-galactosidase. Recruitment of β-arrestin to the GPCR, results in a functional enzyme that generates a chemiluminescent signal when substrate is added. Cells were plated at 5,000 cells per well (40 µL per well) in black walled clear bottomed 384-well plates 24 h prior to running the assay. Medium was removed from the plates. For agonist studies 15 µL of HBSS/HEPES containing compound was added to the cells and the plate was incubated at room temperature for 90 min. For antagonist studies 15 µL of HBSS/HEPES containing compound was added to the cells and was incubated for 15 min prior to the addition of 15 µL of an $EC_{80}$ concentration of agonist. The plate was subsequently incubated at room temperature for 90 min. Both assays were terminated by addition of 15 µL of a Beta-Glo® solution (Promega). Following an additional 30 min incubation the luminescence of each well was measured to determine the level of receptor activation.

**Amine Transporter Assays**

The amine transporter assay measures the ability of compounds to inhibit the activity of the norepinephrine (NET) dopamine (DAT) or serotonin (SERT) transporters by measuring the real time uptake of a dye labeled amine. HBSS/HEPES containing compound (5 µL) was added to the wells of black walled clear bottomed 384-well plate. Transporter dye (25 µL) (Molecular Devices, Sunnyvale CA, USA; Cat # R8174) was added to each well. Finally, 15,000 cells (20 µL) stably expressing the amine transporter of interest were added to each well and the plate is incubated at

37°C for 30 min (DAT) or 60 min (NET and SERT). The plate is transferred to the FLIPR Tetra® instrument and the fluorescence of each well was measured (Ex 470-495 nM; Em 515-575 nM). The level of fluorescence measured directly relates to the level of uptake of the dye labelled amine, with a reduction in levels being related to an inhibition of the respective transporter.

**Phosphodiesterase Assays**

The phosphodiesterase (PDE) assays measure the conversion of 3', 5'-[3H] cAMP to 5'-[3H] AMP (for PDE 3A1 and 4D3) or 3', 5'-[3H] cGMP to 5'-[3H] GMP (for 5A1) by the relevant PDE enzyme subtype. Yttrium silicate (YSi) scintillation proximity (SPA) beads bind selectively to 5'-[3H] AMP or 5'-[3H] GMP, with the magnitude of radioactive counts being directly related to PDE enzymatic activity. The assay was performed in white walled opaque bottom 384-well plates. Test compound (1 µL) in dimethyl sulfoxide was added to each well. Enzyme solution was then added to each well in buffer (in mM: Trizma, 50 (pH7.5); $MgCl_2$, 1.3 mM) containing Brij 35 (0.01% (v/v)). Subsequently, 20 µL of 3',5'-[3H] cGMP (125 nM) or 20 µL of 3',5'-[3H] cAMP (50 nM) was added to each well to start the reaction and the plate was incubated for 30 min at 25°C. The reaction was terminated by the addition of 20 µL of PDE YSi SPA beads (Perkin Elmer, Waltham, MA). Following an additional 8 h incubation period the plates were read on a MicroBeta radioactive plate counter (Perkin Elmer, Waltham, MA, USA) to determine radioactive counts per well.

**Bromodomain-Containing Protein 4 (BRD4) Binding Assay**

The BRD4 fluorescent polarization binding assay uses purified His-tagged BRD4 protein and its interaction with a Cy5 labelled small molecule probe that binds to the BRD4 site involved in the interaction with tetra-acetylated histone H4 peptide. In brief, the assay is performed in low volume black 384 well flat-bottomed polystyrene plates. Compound/vehicle or standard (5 µL) were added to wells followed by His-tagged BRD4 (10 µL; 40 nM final concentration in assay). Following a 15 min incubation at room temperature a proprietary Cy5-labelled probe molecule (5 µL; 2 nM final concentration in assay) was added. Following, an additional 16 h incubation at room temperature fluorescence polarization measurements were made using an Envision plate reader (Perkin Elmer, Waltham, MA, USA) and mP values were used for analysis.

**Acetylcholinesterase Assay**

The assay described is based on Ellmans method, in which thiocholine produced by the action of acetylcholinesterase forms a yellow color with 5,5'-dithiobis(2-nitrobenzoic acid). The intensity of the product color, measured at 405 nm, is proportionate to the enzyme activity in the sample. To each well of a clear 96 polystyrene plate 90 µL enzyme solution (1mU/well) or phosphate buffered saline (PBS) and 10 µL compound/standard or vehicle was added. The plate was incubated at room temperature for 15 min. Subsequently 100 µL of substrate/detection reagent (800 µM acetylthiocholine/1mM 5,5'-dithiobis(2-nitrobenzoic acid)) was added and the plate was read at the 20 min time point.

**hERG Binding Assay**

Human embryonic kidney (HEK) cells stably transfected with a doxycycline inducible plasmid expressing the hERG channel (Accession Number: NM_000238) were cultured in suspension in Ex-cell 293 Serum Free Medium containing fetal bovine serum (5% v/v), L-Glutamine (6 mM), Blasticidin (5 µg/ml) and Zeocin (600 µg/ml) at 37 °C in a humidified environment (5% CO2/95% air). hERG expression was induced by the addition of doxycycline (1 µg/ml) 48 h prior to harvesting by centrifugation. Cell pellets were resuspended in ice cold homogenization buffer (1 mM EDTA, 1 mM EGTA, 1 mM NaHCO3, and cOmplete™ protease Inhibitor cocktail). Cells were homogenized using a dounce homogenizer (20 strokes), and centrifuged (1,000xg) for 10 min at 4ºC. The supernatant was transferred to a new tube and was centrifuged a second time (25,000xg) for 20 minutes at 4ºC. The supernatant was discarded, and the pellet was resuspended in buffer (50 mM HEPES, 10 mM MgCl2, bovine serum albumin (0.2% w/v) and cOmplete™ protease inhibitor cocktail). The samples were adjusted to 5 mg/ml and frozen. For the assay, membrane aliquots were thawed on ice and diluted to 200 µg/ml in assay buffer (25 mM HEPES, 15 mM KCl, 1 mM MgCl2, and 0.05% (v/v) Pluronic F127). A Cy3B tagged N-desmethyl dofetilide ligand was prepared in the same assay buffer solution (5 nM). Compound or vehicle (DMSO) was spotted into each well of a black 384-well low-volume plate. Membrane homogenate (15 µL) and Cy3B tagged ligand (10 µL) were then added to each well and the plate was incubated at room temperature for 16 h. Fluorescence polarization measurements were made using an Envision plate reader (Perkin Elmer, Waltham, MA, USA) and mP values were used for analysis. Binding Ki values were determined using the Cheng-Prusoff equation (Ki = IC50/(1+L/Kd)), where L was the labelled ligand concentration in the assay (2 nM), and the Kd value (1.35 nM) the affinity constant for the labelled ligand.

**hERG, Nav1.5 and Cav1.2 Ion Channel Profiling**

Ionic currents were evaluated in the whole-cell configuration using the Qube384 automated planar patch clamp platform (Sophion Bioscience A/S, Ballerup, Denmark). QChip 384X plates, containing 10 patch clamp holes per well, were used to maximize success rate, which was routinely > 95%.

For hERG experiments, the external solution was composed of (in mM): 132 NaCl, 4 KCl, 1.8 $CaCl_2$, 1.2 $MgCl_2$, 10 HEPES, 11 Glucose, pH 7.4, 305 mOsM. The internal solution contained (in mM): 15 NaCl, 60 KCl, 1 $MgCl_2$, 5 EGTA, 5 HEPES, 70 KF, pH 7.2, 300 mOsM. For Cav1.2 experiments, the external solution was composed of (in mM): 137.9 NaCl, 5.3 KCl, 0.49 $MgCl_2$, 10 $CaCl_2$, 10 HEPES, 0.34 $Na_2HPO_4$, 4.16 $NaHCO_3$, 0.41 $MgSO_4$, 5.5 glucose, pH 7.4, 310 mOsM. The internal solution contained (in mM): 27 CF, 112 CsCl, 2 $MgCl_2$, 10 EGTA, 10 HEPES, 2 $Na_2ATP$, pH 7.2, 305 mOsM. For Nav1.5 experiments (peak current), the external solution was composed of (in mM): 137.9 NaCl, 5.3 KCl, 0.49 $MgCl_2$, 1.8 $CaCl_2$, 10 HEPES, 0.34 $Na_2HPO_4$, 4.16 $NaHCO_3$, 0.41 $MgSO_4$, 5.5 glucose, pH 7.4, and osmolarity of 305 mOsM. The internal solution contained (in mM): 92 CsF, 55 CsCl, 2 $MgCl_2$, 5 EGTA, 5 HEPES, 1 MgATP, pH 7.2, 300 mOsM.

The hERG current was elicited from a holding potential of -80 mV by a voltage step to +40 mV for 500 ms, followed by a repolarizing ramp to -80 mV at -0.6 mV/ms. This pattern was repeated at a rate of 0.05 Hz. Peak hERG current was measured during the ramp. The Cav1.2 current was elicited by a voltage step to 0 mV for 150 ms from a holding potential of -40 mV. Voltage steps were repeated at 0.05 Hz, and Cav1.2 amplitude was measured as the peak current at 0 mV. For Nav1.5 current, from a holding potential of -80 mV, a 200 ms prepulse to -120 mV was used to homogenize channel inactivation, followed by a 40 ms step to a test potential of -15 mV. Membrane potential was further depolarized to +40 mV for 200 ms to completely inactivate the peak Nav1.5 current, followed by a ramp from +40 mV to -80 mV (-1.2 mV/ms). This voltage pattern was repeated at 0.1 Hz, with peak Nav1.5 defined as the maximum current during the step to -15 mV. All studies were conducted at 23° C.

Compounds were dissolved and initially diluted in dimethyl sulfoxide (DMSO), with a final dilution in external solution to generate final working concentrations. The final DMSO concentration in all experiments was 0.33% (v/v).

For all protocols three vehicle periods each lasting 5 minutes were applied to establish a stable baseline. For the standard protocol this was followed by the addition of increasing concentrations of test compound, with each exposure lasting 5 minutes. For the "extended" protocol following the three-vehicle additional each well subsequently received a single concentration of compound. This application was repeated three times for each well, via a flowthrough addition where the solution was replaced with the same compound concentration with each addition. Each exposure lasted 10 minutes.

Patch clamp data were analyzed using Assay Software 6.4.72 (Sophion Bioscience A/S, Ballerup, Denmark). Current amplitudes were determined by averaging the last 4 currents under each test condition. The percentage inhibition of each compound was determined by taking the ratio of current amplitude measured in the presence of various concentrations of the test compound (ICompound) versus the vehicle control current (IVehicle):

% Inhibition = [1-(ICompound/IVehicle)] * 100%.

A dose-response curve was generated and fit to the Hill equation by the Sophion Analyzer software to determine an $IC_{50}$ value for each compound. The minimum and the slope of the fit were free fitted, with the top being fixed to 100% inhibition.

**GABA Patch Clamp Assay**

Compound effects on the human $GABA_A$ receptor (α1β2γ2), stably expressed in human embryonic kidney (HEK) cells, were examined in three modes of action: agonist, antagonist and positive allosteric modulation (PAM) modes. Chloride currents evoked by the activation of the $GABA_A$ receptor were recorded in the whole-cell patch clamp configuration with the automated Qube384® platform (Sophion Bioscience A/S, Baltorpvej, Denmark). The intracellular solution contained (in mM): CsF 90, CsCl 50, $MgCl_2$ 2 EGTA 10, HEPES 10, pH adjusted to 7.2 with CsOH. The extracellular solution contained (in mM): NaCl 138, KCl 5.3, $CaCl_2$ 5, $MgCl_2$ 0.49, HEPES 10, glucose 5.5, $Na_2HPO_4$ 0.34, $NaHCO_3$ 4.16, $MgSO_4$ 0.41, pH adjusted to 7.4 with NaOH. The osmolarity of the internal and external solutions were adjusted with sucrose to 300 mOsm and 305 mOsm, respectively.

Compounds were dissolved and initially diluted in dimethyl sulfoxide (DMSO), with a final dilution in external solution to generate final working concentrations. The final DMSO concentration in all experiments was 0.33% (v/v).

Cells and solutions were loaded into the Qube384 10X Qchip (10 recording wells per well, 384 wells per QChip). After whole-cell configuration was achieved by negative pressure pulse, cells were maintained at a holding potential of -80 mV throughout the experiment. To avoid desensitization and current rundown due to persistent activation of $GABA_A$, a "stacked pipette" approach was utilized to ensure only a brief exposure of the cells to GABA-containing solutions. In this approach, the perfusion pipettes drew from two liquid sources, first from extracellular solution (14 μL), and next from a GABA-containing solution (7 μL). When dispensed into the wells, this resulted in an exposure to GABA lasting 0.8 s, followed by immediate washout by extracellular solution. At the start of the experiment, a baseline $GABA_A$ current was established for each well in response to activation by 40 μM GABA. Each subsequent test condition was normalized against this baseline GABA current on a per well basis.

Agonist effects were measured by recording the current evoked by the test article in the absence of GABA. Agonism (%) was calculated relative to the current produced by 40 μM GABA for each well: [% Agonism = ($I_{\text{test article}}$ / $I_{40\mu\text{M GABA}}$) * 100%].

Antagonist effects of test article were examined in the presence of 40 µM GABA following a 4 min incubation in test article. Antagonism was calculated relative to the current produced by 40 µM GABA for each well [% Antagonism = ($I_{test\ article,\ 40\mu M\ GABA}$ / $I_{40\mu M\ GABA}$) * 100%]. Positive allosteric modulation (%) by test article was detected when current was enhanced relative to the 40 µM GABA normalization current.

$EC_{50}/IC_{50}$ values were calculated by fitting concentration response curve data to a 4-parameter logistic regression equation [% effect=Bottom + (Top-Bottom)/(1+10^((Log$IC_{50}$-Concentration)*HillSlope))].

**Data Analysis**

Agonist/antagonist curves were plotted from individual experiments, and $EC_{50}/IC_{50}$ values were determined using a four-parameter logistic fit. $EC_{50}$ is defined as the concentration of the test article that produced a response that was equal to 50% of the maximal system response. $IC_{50}$ is defined as the concentration of the test article that produced a 50% inhibition of a maximal response. An apparent $K_B$ value for antagonist activity was calculated using the following equation:

Apparent $K_B = IC_{50}/(1+([A]/\text{Agonist } EC_{50})$

where the KB value is the dissociation constant of antagonist for the receptor, $IC_{50}$ is the response produced by the test article in the presence of [A], the concentration of agonist used in the assay. Agonist $EC_{50}$ is the $EC_{50}$ value of the reference agonist used in the assay when tested alone.
